# Identification of a Novel Susceptibility Marker for SARS-CoV-2 Infection in Human Subjects and Risk Mitigation with a Clinically Approved JAK Inhibitor in Human/Mouse Cells

**DOI:** 10.1101/2020.12.09.416586

**Authors:** Marianne R. Spalinger, Rong Hai, Jiang Li, Alina N. Santos, Tara M. Nordgren, Michel L. Tremblay, Lars Eckmann, Elaine Hanson, Michael Scharl, Xiwei Wu, Brigid S. Boland, Declan F. McCole

**Affiliations:** Division of Biomedical Sciences, School of Medicine, University of California Riverside, Riverside, California, USA; Department of Gastroenterology and Hepatology, University Hospital Zurich, and University of Zurich, Zurich, Switzerland; Department of Microbiology & Plant Pathology, University of California Riverside, Riverside, California, USA; Department of Biochemistry and Goodman Cancer Research Centre, McGill University, Montreal, Quebec, Canada; Division of Gastroenterology, University of California San Diego, La Jolla, California, USA; Integrative Genomics Core, Beckman Research Institute of City of Hope, Monrovia, California, USA

**Author notes:** **Corresponding author:** Declan F. McCole, PhD, Division of Biomedical Sciences, School of Medicine, University of California Riverside, 307 School of Medicine Research Building, 900 University Avenue, Riverside, CA 92521. **Author contribution:** MRS: study and experiment design, data acquisition, analysis, and interpretation, writing of the manuscript; RH, MLT: reagent generation, data analysis and critical intellectual input; TMN: data interpretation and critical intellectual input; ANS, JL: data acquisition, sample collection; LE and EH helped develop the mouse model; BSB provided patient serum samples; MS: collection and analysis of patient samples; DFM: study design, supervision of the experiments, data interpretation, funding, writing of the manuscript. All authors reviewed the manuscript.

**Keywords:** Coronavirus, COVID-19, SARS-CoV-2, autoimmune disease, genetic susceptibility, Inflammatory Bowel Disease, Tofacitinib

## Abstract

Coronavirus disease (COVID-19), caused by SARS-CoV-2, has affected over 65 million individuals and killed over 1.5 million persons (December 8, 2020; www.who.int)^1^. While fatality rates are higher among the elderly and those with underlying comorbidities^2^, host factors that promote susceptibility to SARS-CoV-2 infection and severe disease are poorly understood. Although individuals with certain autoimmune/inflammatory disorders show increased susceptibility to viral infections, there is incomplete knowledge of SARS-CoV-2 susceptibility in these diseases.^3–7^ We report that the autoimmune *PTPN2* risk variant rs1893217 promotes expression of the SARS-CoV-2 receptor, ACE2, and increases cellular entry mediated by SARS-CoV-2 spike protein. Elevated ACE2 expression and viral entry were mediated by increased JAK-STAT signalling, and were reversed by the JAK inhibitor, tofacitinib. Collectively, our findings uncover a novel risk biomarker for increased expression of the SARS-CoV-2 receptor and viral entry, and identify a clinically approved therapeutic agent to mitigate this risk.

---

Despite tremendous effort to understand COVID-19 pathogenesis, risk factors for severe disease are still poorly defined. While most attention has focused on symptoms in the airways, gastrointestinal (GI) symptoms were reported in 46% of all cases and 33% presented with GI symptoms in the absence of respiratory symptoms.^8,9^ GI symptoms are associated with longer duration and more severe COVID-19 (e.g. increased prevalence of acute renal insufficiency^10^), emphasizing their importance for early diagnosis and prognosis.^11^ SARS-CoV-2 can directly infect intestinal epithelial cells,^12,13^ and viral particles have been detected in feces even after virus clearance from the respiratory tract^14,15^. This indicates viral shedding in the gut, which may serve as a reservoir of virus replication, and possible oral-fecal transmission although presence of live virus in feces is disputed.^12,16^

SARS-CoV-2 entry into host cells is mediated by its spike glycoprotein (S protein), which is cleaved by cell surface-associated transmembrane protease serine protease 2 (TMPRSS2) and TMPRSS4 to generate the S1 and S2 subunits in a so-called ‘priming’ process.^16,17^ S1 binds to angiotensin I converting enzyme 2 (ACE2), a monocarboxypeptidase controlling cleavage of several peptides within the renin-angiotensin system.^16,17^ S2 drives the subsequent fusion of viral and host membranes.^18^ Interferon (IFN)-JAK-STAT signaling is a suggested major driver of ACE2 expression likely via STAT1/3 binding sites in the ACE2 promoter^19^. ACE2, TMPRSS2, and TMPRSS4 are highly expressed on the surface of epithelial cells such as lung type 2 pneumocytes and absorptive intestinal epithelial cells.^12,18–21^

About 16-20% of the general population carries the single nucleotide polymorphism (SNP) rs1893217 located in the gene locus encoding protein tyrosine phosphatase non-receptor type 2 (PTPN2, also called TCPTP)^22,23^. This SNP causes PTPN2 loss of function and is associated with increased risk for chronic inflammatory and autoimmune diseases including inflammatory bowel disease (IBD), Type 1 diabetes (T1D), and rheumatoid arthritis (RA).^24,25^ PTPN2 directly dephosphorylates several transducers of cytokine receptor signaling including the STAT family of transcription factors (STATs 1/3/5/6)) and Janus kinases (JAK)1 and JAK 3 that are activated by inflammatory cytokines such as IFNγ.^26–28^ JAK inhibitors have emerged as an effective new therapeutic class in many patients with autoimmune diseases. Indeed a JAK-inhibitor, baricitinib, is in clinical trial (ACTT-2) to reduce disease severity and hospitalization time in COVID-19^29^. Tofacitinib (Xeljanz^®^) is a pan-JAK inhibitor that preferentially inhibits JAK1 and JAK3, is approved to treat RA and the IBD subtype, ulcerative colitis (UC), and we have shown that tofacitinib corrects the consequences of PTPN2-loss in IECs^30^.

Using intestinal samples and peripheral blood mononuclear cells (PBMC) from IBD patients harbouring the autoimmune *PTPN2* risk variant rs1893217, human intestinal and lung epithelial cell lines as well as *Ptpn2-*deficient mouse models, we determined that PTPN2-loss promotes ACE2 expression and entry of viral particles expressing SARS-CoV-2 spikes. Elevated ACE2 expression and viral entry were mediated by increased epithelial JAK-STAT signalling, and were reversed by the clinically-approved JAK inhibitor, Tofacitinib. Collectively, our findings not only describe a risk factor for increased SARS-CoV-2 invasion (entry), but also identify a clinically approved drug that may be utilized to mitigate this risk^31^.

## Results

### PTPN2 regulates ACE2 expression *in vivo* and *in vitro*

Mucosal biopsy samples from IBD patients in the Swiss IBD cohort previously genotyped for the IBD-associated loss-of-function SNP *rs1893217* in *PTPN2*^32^ were subjected to RNA sequencing. Using the <u>D</u>atabase for <u>A</u>nnotation, <u>V</u>isualization and <u>I</u>ntegrated <u>D</u>iscovery (DAVID), gene ontogeny (GO) biological pathway analysis indicated “***Digestion***” as the most significantly regulated function based on *PTPN2* genotype (presence of risk ‘C’ allele, patients with the CT or the CC genotype) independent of disease severity (Supplementary Table 1). This biological process included the *ACE2* gene, which was also found in three other processes that were increased in ‘C’ allele carriers (Supplementary Table 1). Quantitative PCR and Western blotting on intestinal biopsies isolated from Crohn’s disease (CD) and ulcerative colitis (UC) patients (Supplementary Table 2) confirmed increased mRNA and protein expression of ACE2 in C-allele carriers (Figure 1A-B). Furthermore ACE2 protein expression negatively correlated with PTPN2 phosphatase activity (Figure 1C). To confirm these findings in *Ptpn2-*knockout (KO) mice, which exhibit a severe inflammatory phenotype and die within few weeks after birth^33^, we explored ACE2 expression in 3-week-old mice when the intestinal epithelium appears relatively normal compared to heterozygous (Het) and wild-type (WT) littermates. Although *Ptpn2*-KO mice did not exhibit any difference in *Ace2* mRNA expression in whole intestinal samples compared with wild-type (WT) mice (Supplementary Figure 1A), *Ace2* mRNA and protein expression in intestinal epithelial cells (IEC) from these mice was significantly increased (Supplementary Figure 1A, Figure 1D). In addition, *Ace2* mRNA expression in lung and cardiac tissue was significantly increased in *Ptpn2*-KO mice (Figure 1D, Supplementary Figure 1B-C). Given the strong increase of ACE2 expression in IECs of *Ptpn2*-KO mice, and to explore whether PTPN2 regulates ACE2 in the absence of inflammation, we confirmed these findings in mice lacking PTPN2 specifically in IECs (*Ptpn2*^ΔIEC^ mice). Also in these mice, *Ace2* mRNA and protein expression were clearly elevated in PTPN2-deficient IECs (Figure 1E, Supplementary Figure 1D), demonstrating that the increase in *Ace2* expression was not dependent on inflammation. This effect was confirmed in intestinal epithelial, lung epithelial and monocyte cell lines upon *PTPN2* knockdown, where depletion of *PTPN2* resulted in elevated *ACE2* expression (Supplementary Figure 1E).^28^ Notably, the serine proteases TMPRSS2 and TMPRSS4, which are additional cofactors of SARS-CoV-2 viral entry, were not altered in PTPN2 knockdown (PTPN2-KD) cells (Supplementary Figure 2). This suggests that PTPN2 specifically regulates *ACE2* expression.

**Figure 1.**
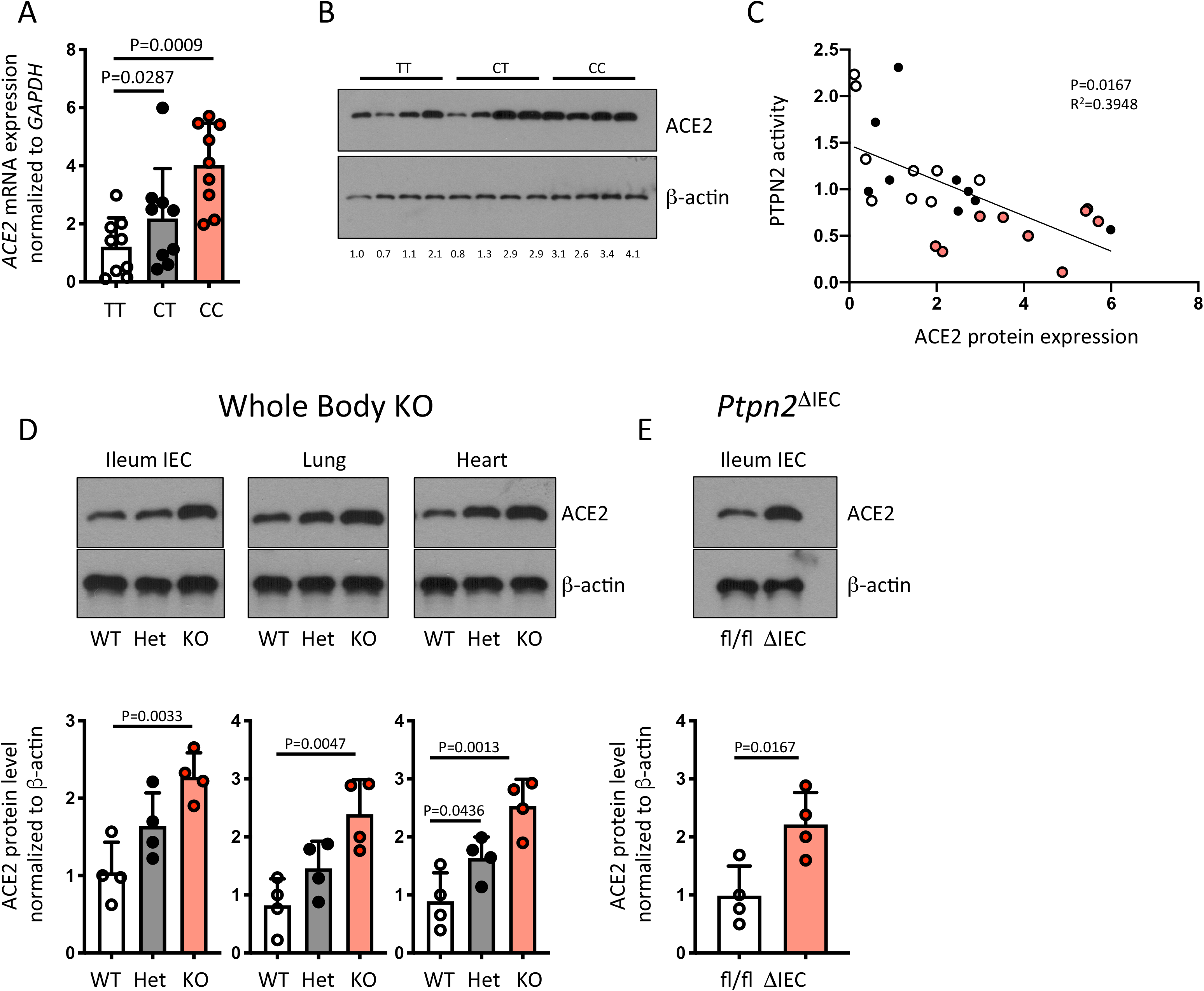
Presence of SNP rs1893217 in ***PTPN2*** promotes ACE2 expression. **A)** Ileum and colon biopsies from IBD patients homozygous for the major allele (TT), heterozygous (CT) or homozygous for the inflammatory disease-associated minor allele (CC) in *PTPN2* SNP rs1893217 were analyzed for **A)** *ACE2* mRNA and **B)** ACE2 protein expression. Depicted are representative Western blot pictures and values below the blot indicate relative band density normalized to β-actin and TT controls. **C)** PTPN2 phosphatase activity levels were analyzed in the same samples as in B and correlated with relative PTPN2 protein levels. D+E) Representative Western blot pictures and respective densitometric analysis for ACE2 and β-actin in intestinal epithelial cells (IEC) from the illeum, lung tissue, and heart tissue from 3-week-old wild-type (WT), whole-body *Ptpn2* heterozygous (Het) or *Ptpn2* knock-out (KO) mice (D) or mice in which *Ptpn2* was specifically deleted in IECs (ΔIEC) or their control littermates (fl/fl) **(E)**. Statistical differences are indicated in the figures (One-way ANOVA (A+B, D+E) or linear regression (C)), **A-C**: n = 8 per genotype, D+E: n = 4. Each dot represents a biological replicate.

### PTPN2 negatively regulates SARS-CoV-2 spike protein-mediated viral entry into epithelial and immune cells

To assess the functional consequence of increased ACE2 expression in PTPN2-deficient cells, we assessed whether deletion of PTPN2 affects the uptake of virus-like particles (VLPs) expressing SARS-CoV-2 spikes. Apical uptake of empty VLPs (-), which served as a negative control, or uptake of VLPs covered with spike G glycoprotein of the rhabdovirus vesicular stomatitis virus (G), which served as a positive control for viral entry, was not affected in PTPN2-KD Caco-2BBe (Figure 2A) and HT-29.cl19A IECs (Figure 2B) or A549 lung epithelial cells (Figure 2C), thus indicating that non-specific uptake was not affected upon PTPN2 knockdown. In contrast, VLPs expressing SARS CoV-2 spikes entered PTPN2-KD epithelial cells more efficiently than control cells (Figure 2A-C), indicating that PTPN2 deficiency not only promotes ACE2 expression but also viral uptake into intestinal and lung epithelial cells. Notably, loss of PTPN2 also caused a significant increase in ACE2 expression in PTPN2-KD monocytes (Supplementary Figure 1) and increased CoVS entry (Figure 2D). This indicates that PTPN2 regulation of ACE2 and SARS-CoV-2 spike-expressing VLP entry is not restricted to epithelial cells but has similar functional consequences in immune cells. Furthermore, increased uptake of SARS-CoV-2 spike-expressing VLPs in PTPN2-deficient cells was no longer observed upon inhibition of ACE2 with a blocking antibody (Supplementary Figure 3), indicating that ACE2 mediated SARS-CoV-2 spike-expressing VLP uptake. In summary, this indicates that loss of PTPN2 promotes SARS-CoV-2 uptake by promoting ACE2 expression.

**Figure 2.**
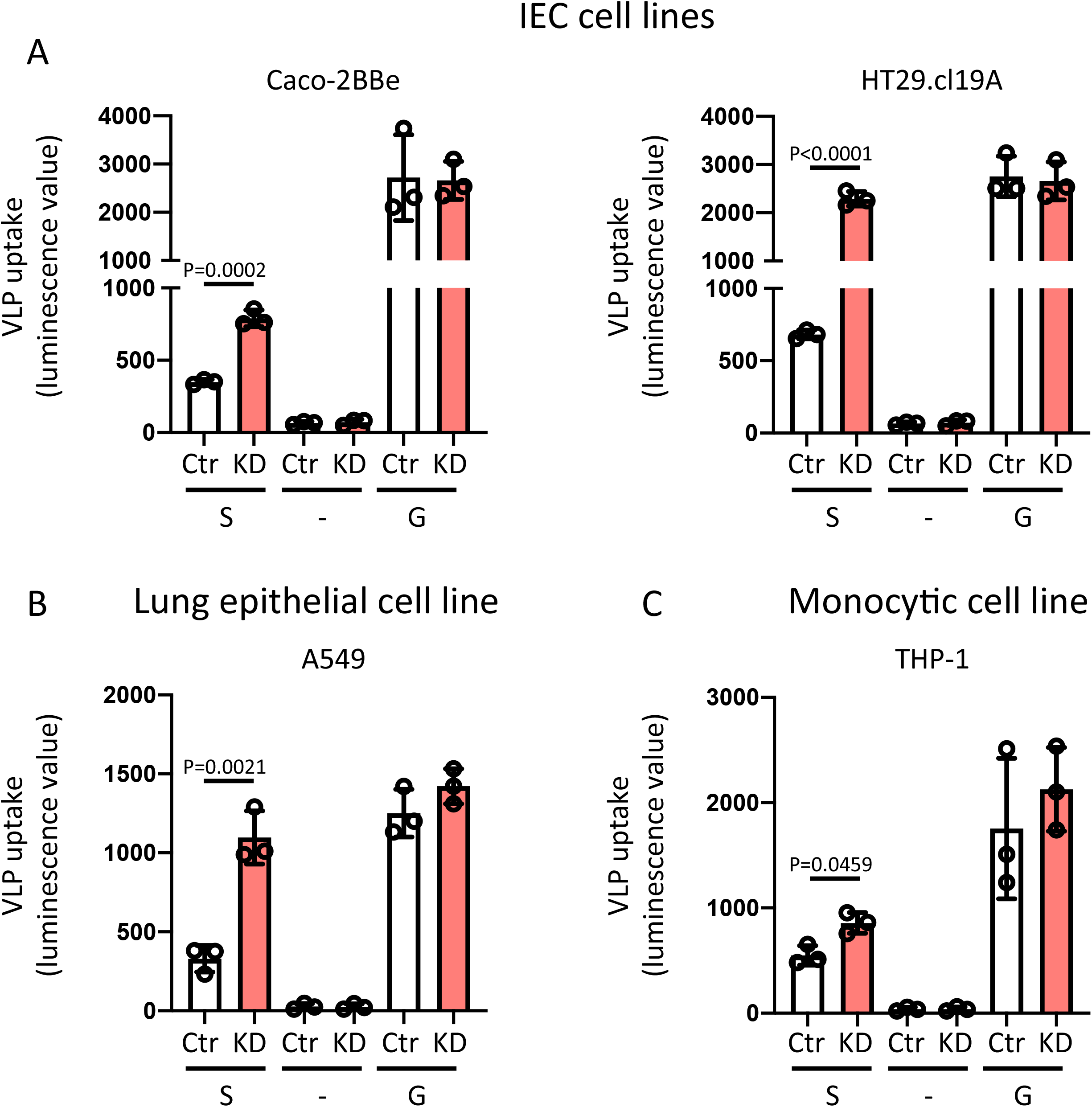
PTPN2 knockdown facilitates entry of VLPs expressing SARS-CoV-2 spike S protein. **A)** Caco-2BBe, **B)** HT-29.cl19A, **C)** A549, and **D)** THP-1 cells expressing non-targeting control (Ctr) or *PTPN2-*specific (KD) shRNA were incubated with virus like particles (VLPs) expressing renilla luciferase and SARS-CoV-2 spike protein (S), no additional proteins (-, negative control), or the spike G glycoprotein of the rhabdovirus vesicular stomatitis virus (G, positive control). 48 h after infection, luminescence values of the supernatant were measured as an approximation of VLP uptake. Statistical differences are indicated in the figure (Student’s t test, n = 3 independent experiments). Each dot represents the average of an independent experiment with 2-3 technical replicates, each.

### IFN-α promotes ACE2 expression and SARS-CoV-2 Spike protein uptake

It has been suggested that ACE2 expression is induced by interferons^34^ and its promoter is reported to have putative STAT1 binding sites^19,35^, although newer findings debate whether interferons can induce ACE2 expression, or if it drives expression and release of a shorter version of the protein^36^. Since PTPN2 is a potent suppressor of IFN-γ-induced signaling cascades^27,37^, and directly dephosphorylates STAT1^38^, we assessed whether IFN-γ treatment promotes ACE2 expression in our cell culture models. Indeed, IFN-γ promoted ACE2 expression and this was further increased in PTPN2-KD cells (Figure 3A, Supplementary Figure 4). Silencing of STAT1 using siRNA constructs prevented IFN-γ-induced ACE2 mRNA and protein expression and STAT1-siRNA treated PTPN2-KD Caco-2BBe, A549 and THP-1 cells expressed ACE2 levels comparable to those in Ctr cells (Figure 3B+C and Supplementary Figure 5). In line with these effects on ACE2 expression, uptake of SARS-CoV-2 S protein-expressing VLPs was elevated in IFN-γ-treated control and PTPN2-KD cells, while STAT1 silencing normalized the uptake, both in Ctr cells and PTPN2-KD cells (Figure 3D+E, Supplementary Figure 5). This strongly suggests that deletion of PTPN2 promotes ACE2 expression and SARS-CoV-2 entry in a STAT1-dependent manner and inhibition of STAT signaling may mitigate elevated ACE2 expression and SARS-CoV-2 entry into host cells.

**Figure 3.**
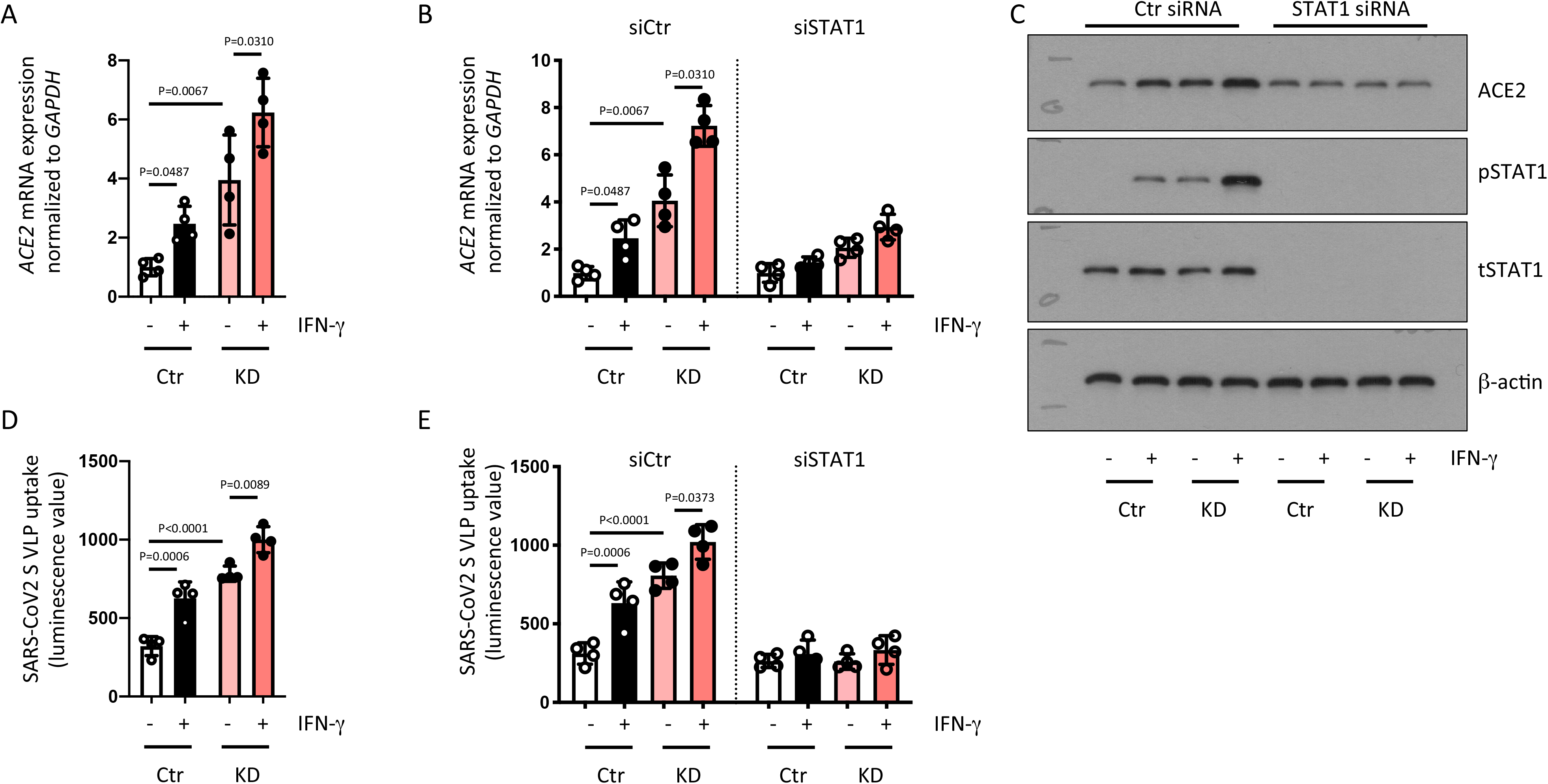
IFN-α promotes uptake of SARS-CoV2 spike-expressing VLPs in a STAT1-dependent manner. **A)** Caco-2BBe cells expressing non-targeting control (Ctr) or *PTPN2-*specific (KD) shRNA were treated with IFN-α for 24 h and analyzed for *ACE2* mRNA expression. **B)** Ctr and KD Caco-2BBe cells were treated with non-targeting control (siCtr) or STAT1-specific (siSTAT1) siRNA prior to incubation with IFN-α for 24 h and analysis for *ACE2* mRNA expression. **C)** Representative Western blot images for the indicated proteins from cells treated as in B. **D)** Ctr and KD Caco-2BBe cells were infected with VLPs expressing SARS-CoV-2 spike protein in the presence or absence of IFN-α and luminescence measured after 48 h. **E)** Ctr and KD Caco-2BBe cells were treated with non-targeting control (siCtr) or STAT1-specific (siSTAT1) siRNA prior to infection with VLPs expressing SARS-CoV-2 spike protein in the presence or absence of IFN-α and luminescence measured after 48 h. Statistical differences are indicated in the figure (One-way ANOVA, n = 4). Each dot represents the average of an independent experiment with 2-3 technical replicates, each.

### Tofacitnib reverses ACE2 overexpression and increased SARS-CoV-2 entry in PTPN2-deficient human cells and mice

To test whether inhibition of STAT activation can indeed prevent elevated ACE2 expression in PTPN2-deficient cells, we next treated Caco-2BBe IECs with the pan-JAK-inhibitor tofacitinib. Similar to our findings in cells treated with STAT1 siRNA, inhibition of JAK-STAT signalling in Caco-2BBe cells using tofacitinib prevented IFN-γ-induced increases in ACE2 mRNA and protein expression and normalized the elevated ACE2 levels in PTPN2-KD cells (Figure 4A+B). Moreover, tofacitinib reduced ACE2-mediated SARS-CoV-2 spike-expressing VLP uptake in both PTPN2-KD and in IFN-γ-treated cells (Figure 4C). Similar effects were observed in A549 lung epithelial cells (Supplementary Figure 6), indicating that tofacitinib treatment might not only reduce ACE2-mediated intestinal viral uptake, but also reduce viral uptake in the respiratory tract, the primary entry site of SARS-CoV-2. We next tested if tofacitinib altered ACE2 levels in human subjects. Levels of soluble ACE2, which has been suggested to reduce viral binding to host cells^39^, were not altered in UC patients treated with tofacitinib when compared to UC patients under anti-TNF treatment with similar disease activity (Figure 4D, Supplementary Table 3). This suggests that tofacitinib treatment does not reduce shedding/release of ACE2 into serum. In contrast, when analysing ACE2 levels in PBMCs from IBD patients carrying the *PTPN2* loss-of-function SNP rs1893217, we again observed that ACE2 mRNA and protein levels and SARS-CoV-2 spike-expressing VLP entry were clearly elevated in variant carriers compared to non-carriers (Figure 4E-G). Notably, treatment with tofacitinib not only reduced ACE2 levels and viral entry in variant carriers, but also in non-carriers (Figure 4E-G). These findings indicate that loss of PTPN2 or presence of the loss-of-function variant in *PTPN2* promotes ACE2 expression and subsequently facilitates uptake of SARS-CoV-2-spike-expressing VLPs, and that treatment with tofacitinib can mitigate this potential risk by reducing ACE2 cellular expression rather than affecting release of soluble ACE2. Furthermore, our data strongly indicate that treatment with tofacitinib might not only be beneficial in *PTPN2* variant carriers, but also for non-carriers.

**Figure 4.**
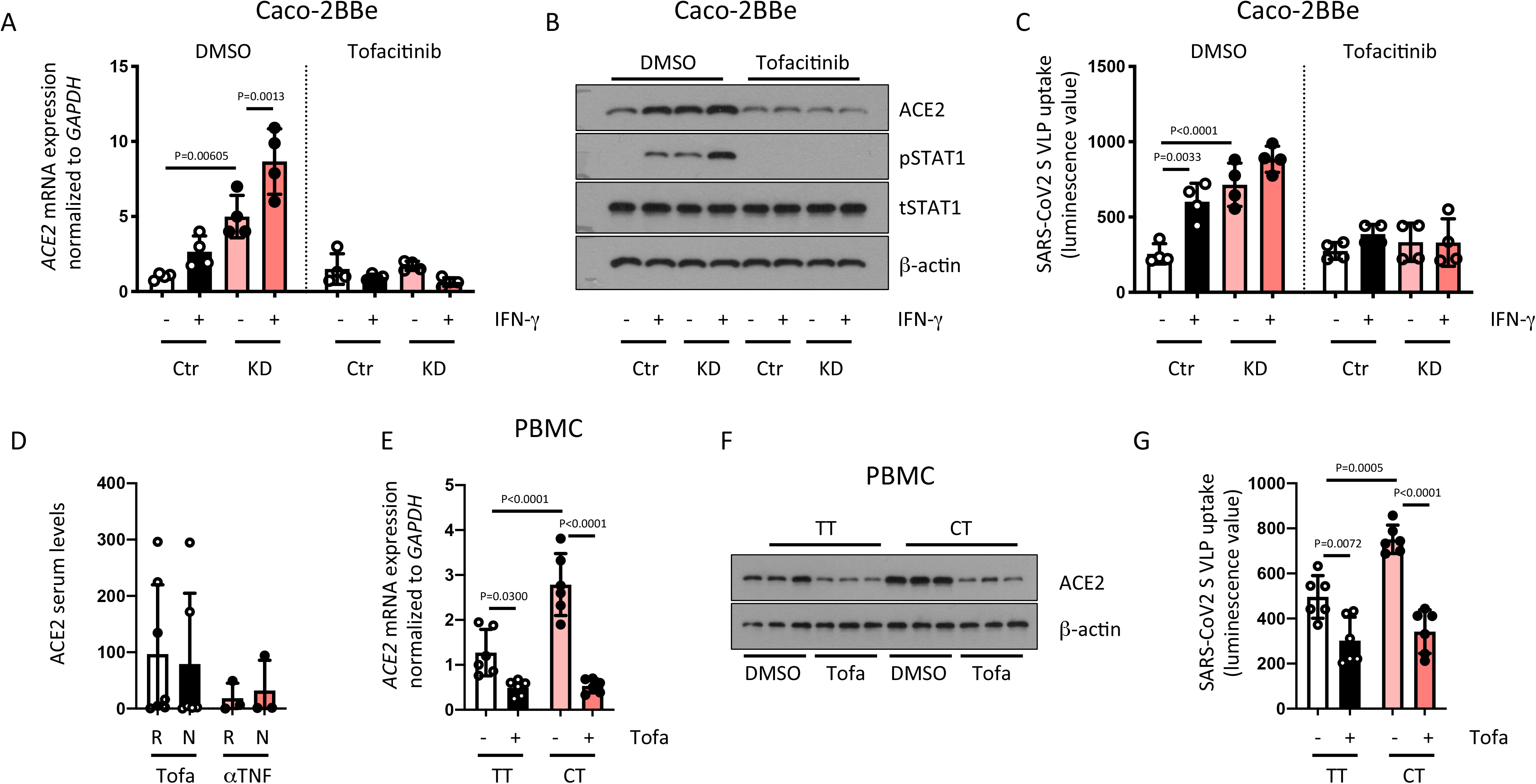
Tofacitinib prevents ACE2 upregulation and reduces SARS-CoV-2 VLP-uptake. **A-C:** Caco-2BBe cells expressing non-targeting control (Ctr) or *PTPN2-*specific (KD) shRNA were treated with vehicle (DMSO) or Tofacitinib for 1 h prior to infection with VLPs expressing SARS-CoV-2 spike protein in the presence or absence of IFN-α. **A)** Relative mRNA expression of *ACE2* and **B)** representative Western blot pictures for the indicated proteins 24 h after VLP treatment, **C)** luminescence values as an approximation of VLP uptake after 48 h. **D)** Serum from ulcerative colitis patients treated with tofacitinib or anti-TNFα were analyzed for ACE2 protein level (R = responder; NR = non-responder). **E-G)** Peripheral blood mononuclear cells from IBD patients homozygous for the major allele (TT) or heterozygous for the disease-associated risk allele in SNP rs1893217 in *PTPN2* were treated with Vehicle (DMSO) or Tofacitinab for 24 h and analyzed for **E)** ACE2 mRNA and **F)** ACE2 protein expression. **G)** After 24 h Tofacitinib-treatment, the cells were infected with VLPs expressing SARS-CoV-2 spike protein and luminescence analyzed after 48 h as an approximation for VLP uptake. Statistical differences are indicated in the figure (One-way ANOVA, A-C: n = 4; D: Tofacitinib treated patients n = 12, anti-TNF-treated patients n = 6); E-G: n = 6). Each dot represents the average of an independent experiment (A-C) or a biological sample (D-G) with 2-3 technical replicates, each.

Our data consistently demonstrate that PTPN2 dysfunction promotes expression of ACE2 and uptake of SARS-CoV-2 spike protein, and this is further increased by inflammation. This is striking given a recent paper identifying that non-genotyped IBD patients (+/− inflammation) showed no change in ACE2 or TMPRSS2 expression, while experimental colitis in mice reduced gut epithelial *Ace2* expression.^40^ This indicates that inflammation does not promote ACE2 expression *per se,* but that the elevated ACE2 levels in patients or mice with reduced PTPN2 activity are indeed due to PTPN2 deficiency.

Summarized, we demonstrate that SNP rs1893217 in *PTPN2* is associated with increased expression of ACE2 and SARS-CoV-2 entry, and potentially represents one of the first identified COVID-19 genetic susceptibility biomarkers. By using samples collected well before the COVID-19 outbreak, our identification of a genetic susceptibility marker avoids the potential for ascertainment bias in most genetic studies of COVID-19, as clinically significant COVID-19 patients are more likely to be included in research projects than asymptomatic cases^41^. With respect to genetic markers of COVID-19 susceptibility, studies have proposed the involvement of ABO blood groups, with blood group O associated with lower risk, while blood group A was associated with higher risk of acquiring COVID-19 compared with non-A blood groups^41–44^. However, this correlation did not culminate in therapeutic implications. Moreover, a cluster of genes on chromosome 3 has been linked with increased severity, although this may have distinct geographic distributions^41,45^. In contrast, our finding of increased ACE2 expression/viral particle uptake in *PTPN2* variant cells might not only indicate a potentially novel genetic marker for increased disease, but also identifies tofacitinib – a drug already approved for treatment of arthritis and IBD – and potentially other JAK inhibitors such as baricitinib, as a potential therapeutic strategy to specifically mitigate this risk.

## Methods

### Patient samples

Samples from IBD patients used for RNA and protein isolation were obtained from the Swiss IBD cohort and the sample use approved by the local ethic’s board (Ethic’s board of the Kanton Zurich, Switzerland; approval number EK1977). Serum samples were obtained from University of California San Diego under IRB Protocol # 131487. All patients provided informed consent.

### Mice

PTPN2-knock-out (KO) mice in the BALB/c background were a gift from Prof. M. Tremblay at McGill University in Montreal. To generate mice lacking PTPN2 specifically in intestinal epithelial cells (ΔIEC mice), mice with a loxP-flanked exon 3 of the PTPN2 gene (PTPN2-fl/fl mice, originally obtained from EUCOMM, abbreviated as fl/fl) were crossed with VillinCre-ERT2 mice (Jackson Laboratories). Translocation of the Cre-ERT2 construct and subsequent recombination and deletion of the floxed gene was induced by tamoxifen-injections (i.p., 1mg/mouse/day) on 5 consecutive days. All mouse experiments were conducted according to protocols approved by the IACUC commission of the University of California Riverside (AUP20190032).

### Protein isolation and Western blotting

Protein isolation and Western blotting were performed according to standard procedures. For protein isolation from cells, the cells were washed with ice cold PBS and lysed in RIPA buffer containing phosphatase and protease inhibitors (Roche, South San Francisco, CA). For mouse and human biopsies, the samples were dissociated in RIPA buffer using a bead-beater and metal beads. All samples were then sonicated for 30 seconds, centrifuged (10 min. at 12’000 g at 4°C), and the supernatant transferred into fresh tubes. Protein concentration was detected using a BCA assay (Thermo Fisher Scientific, Waltham, MA). For Western blots, aliquots with equal amounts of protein were separated by electrophorese on polyacrylamide gels, and the proteins blotted on nitrocellulose membranes. The membranes were then blocked in blocking buffer (3% milk, 1% BSA in tris-buffered saline with 0.5% Tween) for 1 h and incubated over night at 4°C with anti-ACE2 (Clone E-11, Santa Cruz Biotechnology), anti-phospho-STAT1 (Tyr701, clone 58D6; Cell Signalling Technologies, Danvers, MA), anti-STAT1 (clone 42H3; Cell Signalling Technologies), or anti-β-actin (Clone AC-74, Sigma-Aldrich, St. Louis, MO) antibodies. On the next day, the membranes were washed in tris-buffered saline with 0.5% Tween, incubated with HRP-coupled secondary antibodies (Jackson Immunolabs, West Grove, PA), washed again, and immunoreactive proteins detected using ELC substrate (Thermo Fisher Scientific) and X-ray films (GE Healthcare, Chicago, IL).

### PTPN2 phosphatase assay

For PTPN2 activity measurements, 100 μg protein lysates were pre-cleared for 1 h using Sepharose A beads, incubated with 2 μl anti-PTPN2 antibody (Clone D7T7D, Cell Signaling Technologies) over night, incubated with Sepharose A beads for 1 h and centrifuged (3 min. at 12’000 g at 4°C). The precipitates were washed 3 times with ice cold PBS and the beads resuspended in phosphatase assay buffer (Thermo Fisher Scientific) and phosphatase activity measured using the EnzCheck Phosphatase assay (Thermo Fisher Scientific) according to the manufacturer’s instructions.

### RNA isolation and qPCR

For RNA isolation, biopsies were disrupted in RLT buffer (Qiagen, Valencia, CA) using a bead beater and metal beads. Cells were washed twice in ice-cold PBS before lysis in RLT buffer. All samples were then passed 3-5 times through a 26G needle prior to RNA isolation using the RNAeasy mini kit from Qiagen. RNA concentrations were estimated by absorbance measurement at 260 and 280 nm, and cDNA generated using the qScript reverse transcriptase (Quantabio, Beverly MA). Quantitative real-time PCR was performed using iQ SYBR Green Supermix (BioRad, Hercules, CA) on a C1000 Thermal cycler equipped with a CFX96 Real-Time PCR system using BioRad CFX Manager 3.1 Software. Measurements were performed in triplicates using *GAPDH* as an endogenous control. Results were analyzed by the ΔΔCT method. The real-time PCR included an initial enzyme activation step (3 minutes, 95 °C) followed by 45 cycles consisting of a denaturing (95 °C, 10 seconds), an annealing (53°-60°C, 10 seconds) and an extending (72 °C, 10 seconds) step. The used primers are listed in Supplementary Table S3.

### VLPs and measurement of VLP uptake

To produce pseudoparticles, 293T cells were transfected with plasmids encoding a minimal HIV (pTRIP, CSGW, CSPW) provirus expressing the Gaussia Luciferase (Gluc), gag-pol, and S protein of SARS-CoV-2 virus using polyethylenimine (PEI) transfection reagent^46–48^. Supernatants were collected at 24, 48 and 72 hours post-transfection, pooled, filtered (0.45 mm), aliquoted and stored at −80°C. Pseudoparticle infections were performed in the presence of 4 µg/ml polybrene. Appropriate amounts of pseudoparticle were added onto target cells and plates incubated for 3 hrs (37°C) before changing media. Gaussian luciferase was measured at 24, 48, and 72 hrs after infection.

To measure VLP uptake into cells, the VLP-containing medium was diluted 1:2 in cell culture medium and applied to the cells. After 1 h, the cells were washed with PBS and fresh medium added to the cells. In experiments with IFN-γ (1000 IU/ml; Roche, Belmont CA), the replacement medium for cells infected in presence of IFN-γ contained IFN-α as well. To determine VLP uptake, luciferase luminescence in cell culture supernatant was determined using the Renilla luciferase activity assay from Thermo Fisher Scientific.

### Cell culture, PTPN2 knockdown, siRNA treatment and IFN-γ treatment

HT-29.cl19A were obtained from Kim E. Barrett (University of California, San Diego, California), Caco-2BBe, A549 and THP-1 cells were originally obtained from ATCC and cultured according to the manufacturer’s recommendation in medium with 10% FCS. For PTPN2 knockdown, the cells were infected with lentiviral particles containing non-targeting control shRNA (Ctr) or PTPN2-specfic shRNA as described previously^49^ and stable clones selected using puromycin. For STAT1 silencing, the cells were transfected with previously validated, STAT1-specific or non-targeting control siRNA constructs (Dharmacon) using DharmaFECT transfectionre agents as described previously^50^. In experiments with STAT1 siRNA and IFN-γ treatment, the culture medium was replaced with serum-free medium 8 h prior to addition of IFN-γ (1000 IU; Roche). In experiments with Tofacitinib, the cells were treated with tofacitinib (50 µM, MedChemExpress, Monmouth Junction, NJ). Control cells were treated with an equal amount of vehicle (dimethyl sulfoxide, DMSO, 0.5%, Sigma-Aldrich).

### ELISA

Human ACE2 ELISA was obtained from R&D and performed according to the manufacturer’s instructions with undiluted serum.

### RNA sequencing

RNA sequencing was performed and analyzed by the Integrative Genomics Core, City of Hope National Medical Center (Duarte USA). The RNA-seq libraries were constructed with Kapa mRNA HyperPrep Kit (Roche) following the manufacturer’s recommendation. The libraries were then sequenced on an Illumina Hiseq 2500 with single end 50 bp reads to a depth of about 35M. The sequences were aligned to human genome assembly hg38 using Tophat2 v2.0.14. RNA-seq data quality was evaluated using RSeQC v2.5. For each sample, expression counts for Ensembl genes (v92) were summarized by HTseq v0.6.1, and reads per kilobase of transcript per million mapped reads (RPKM) were calculated. Count normalization and differential expression analyses between groups were conducted using Bioconductor package “DESeq2” v1.14.1. Heatmaps were generated using R package “heatmap3”. The Gene Ontology and pathway analysis was performed using DAVID online tools and Ingenuity Pathway Analysis (IPA).

### Statistics

Data are represented as mean of a series of n biological repetitions ± standard deviation (SD). Data followed a Gaussian distribution and variation was similar between groups for conditions analyzed together. Significant differences were determined using GraphPad Prism v9 software using analysis of variance (ANOVA). p-values below 0.05 were considered significant. Mice for ex vivo analyses were matched for age and sex. Numbers of replicates are given in the figure legends. No data points were excluded from statistical analysis.

## Author contribution

MRS: data acquisition, analysis and interpretation, study design, writing the manuscript; JL, ANS: data acquisition; RH, MLT: generation of material for the study and technical input; RH, TMN: critical intellectual input; XW: sequencing and bioinformatics; LE, EH, BSB, MS: human sample acquisition; DFM: conceived the study, data interpretation, manuscript writing, acquisition of funding.

## Additional Information

Supplementary Information is available for this paper.

Correspondence and requests for materials should be addressed to Declan F. McCole.

