## Supplementary tables 1-3, figures 1-6 for "Identification of a Novel Susceptibility Marker for SARS-CoV-2 Infection in Human Subjects and Risk Mitigation with a Clinically Approved JAK Inhibitor in Human/Mouse Cells"

### Supplementary Table 1. Biological processes upregulated in IBD patients with the *PTPN2* rs1893217 variant.

| Category | ID | Term | % | P-Value | Genes | Benjamini |
|---|---|---|---|---|---|---|
| **GOTERM_BP_FAT** | GO:0007586 | Digestion | 6.67 | 1.21E-04 | AKR1C2, MUC3A, SLC5A1, **ACE2**, PRSS1, FABP2 | 7.40E-02 |
| **GOTERM_BP_FAT** | GO:0022600 | Digestive system process | 3.33 | 1.40E-02 | MUC3A, SLC5A1, FABP2 | 9.89E-01 |
| **GOTERM_BP_FAT** | GO:0055114 | Oxidation reduction | 10 | 1.98E-02 | AKR1C3, ALDH1A1, AKR7L, AKR1C2, CYP3A5, CYP2C19, CYP2C18, CYP2C8, CYBRD1 | 9.86E-01 |
| **GOTERM_BP_FAT** | GO:0055085 | Transmembrane transport | 8.89 | 3.13E-02 | SLC5A1, KCNH6, SLC5A9, MFSD2A, ABCC2, SLC46A3, ABCC8, FLVCR1 | 9.94E-01 |
| **GOTERM_BP_FAT** | GO:0001991 | Regulation of systemic arterial blood pressure by circulatory renin-angiotensin | 2.22 | 4.18E-02 | **ACE2**, PCSK5 | 9.96E-01 |
| **GOTERM_BP_FAT** | GO:0010817 | Regulation of hormone levels | 4.44 | 4.65E-02 | SHBG, **ACE2**, PCSK5, SMPD3 | 9.94E-01 |
| **GOTERM_BP_FAT** | GO:0003081 | Regulation of systemic arterial | 2.22 | 5.70E-02 | **ACE2**, PCSK5 | 0.995182193 |

| | | | | | | |
|---|---|---|---|---|---|---|
| | | blood pressure by renin-angiotensin | | | | |
| **GOTERM_BP_FAT** | GO:0050892 | Intestinal absorption | 2.22 | 8.19E-02 | SLC5A1, FABP2 | 0.998877071 |
| **GOTERM_BP_FAT** | GO:0043043 | Peptide biosynthetic process | 2.22 | 8.68E-02 | GGT2, PCSK5 | 0.998362738 |

**Supplementary Table 2. Characteristics of *PTPN2* rs1893217 variant genotyped IBD patients with Crohn's disease (CD) or ulcerative colitis (UC).**

| Genotype | Disease | Severity | location | inflamed | Gender (M/F) | type of sample |
|---|---|---|---|---|---|---|
| TT | UC | quiescent | ileum | no | M | RNA |
| TT | CD | moderate | ileum | yes | M | RNA |
| TT | CD | quiescent | rectum | no | M | RNA |
| TT | UC | moderate | rectum | yes | M | RNA |
| TT | UC | moderate | ileum | yes | F | RNA, Protein, PBMC |
| TT | UC | quiescent | ileum | no | F | RNA, Protein, PBMC |
| TT | CD | severe | rectum | yes | F | RNA, Protein, PBMC |
| TT | CD | moderate | rectum | yes | M | RNA, Protein, PBMC |
| TT | CD | quiescent | rectum | no | M | RNA, Protein, PBMC |
| TT | UC | moderate | ileum | yes | M | Protein, PBMC |
| TT | UC | quiescent | ileum | no | M | Protein |
| TT | UC | severe | rectum | yes | F | Protein |
| TT | CD | quiescent | ileum | no | F | Protein |
| CT | UC | quiescent | ileum | no | M | RNA |
| CT | CD | moderate | ileum | yes | M | RNA |
| CT | CD | quiescent | rectum | no | M | RNA |
| CT | UC | moderate | rectum | yes | M | RNA |
| CT | UC | moderate | ileum | yes | F | RNA, Protein, PBMC |
| CT | UC | quiescent | ileum | no | F | RNA, Protein, PBMC |
| CT | CD | severe | rectum | yes | F | RNA, Protein, PBMC |
| CT | CD | moderate | rectum | yes | M | RNA, Protein, PBMC |
| CT | CD | quiescent | rectum | no | M | RNA, Protein, PBMC |
| CT | UC | moderate | ileum | yes | M | Protein, PBMC |
| CT | UC | quiescent | ileum | no | M | Protein |
| CT | UC | severe | rectum | yes | F | Protein |
| CT | CD | quiescent | ileum | no | F | Protein |
| CC | UC | quiescent | ileum | no | M | RNA |
| CC | CD | severe | ileum | yes | M | RNA |
| CC | CD | quiescent | rectum | no | M | RNA |
| CC | UC | moderate | rectum | yes | M | RNA |
| CC | UC | moderate | ileum | yes | F | RNA+Protein |
| CC | UC | quiescent | ileum | no | F | RNA+Protein |
| CC | CD | severe | rectum | yes | F | RNA+Protein |
| CC | CD | moderate | rectum | yes | M | RNA+Protein |
| CC | CD | quiescent | rectum | no | M | RNA+Protein |
| CC | UC | moderate | ileum | yes | M | Protein |
| CC | UC | quiescent | ileum | no | M | Protein |
| CC | UC | severe | rectum | yes | F | Protein |
| CC | CD | quiescent | ileum | no | F | Protein |

**Supplementary Table 3. Characteristics of ulcerative colitis patients from which serum samples were obtained.**

| Disease | Gender (M/F) | Medication | Anti-TNF naïve | Response anti-TNF | Tofacitinib response | Mayo Score | Endoscopic remission | Clinical response |
|---|---|---|---|---|---|---|---|---|
| UC | M | Tofacitinib | no | no | yes | 0 | yes | yes |
| UC | M | Tofacitinib | no | yes | yes | 1 | yes | yes |
| UC | M | Tofacitinib | no | no | yes | 1 | yes | yes |
| UC | M | Tofacitinib | no | no | yes | 1 | yes | yes |
| UC | M | Tofacitinib | no | no | yes | 0 | yes | yes |
| UC | M | Tofacitinib | no | no | yes | 0 | yes | yes |
| UC | M | Tofacitinib | yes | na | yes | 0 | yes | yes |
| UC | M | Tofacitinib | no | no | yes | 1 | yes | yes |
| UC | M | Tofacitinib | no | no | no | 2 | no | no |
| UC | M | Tofacitinib | no | no | no | 3 | n | no |
| UC | F | Tofacitinib | no | no | no | 3 | no | no |
| UC | M | Tofacitinib | no | no | no | 3 | no | no |
| UC | M | Tofacitinib | no | no | no | 3 | no | no |
| UC | M | infliximab | no | yes | no | 0 | yes | yes |
| UC | M | adalimumab | no | yes | no | 0 | yes | yes |
| UC | M | infliximab | no | yes | no | 0 | yes | yes |
| UC | M | adalimumab | no | no | no | 3 | no | no |
| UC | M | adalimumab | no | no | no | 3 | no | no |
| UC | M | infliximab | no | partial | no | 2 | no | no |

**Endoscopic remission defined as Mayo endoscopic score of 0 or 1**

**Clinical response defined as >50% reduction in clinical symptoms**

SUPPLEMENTARY FIGURE 1

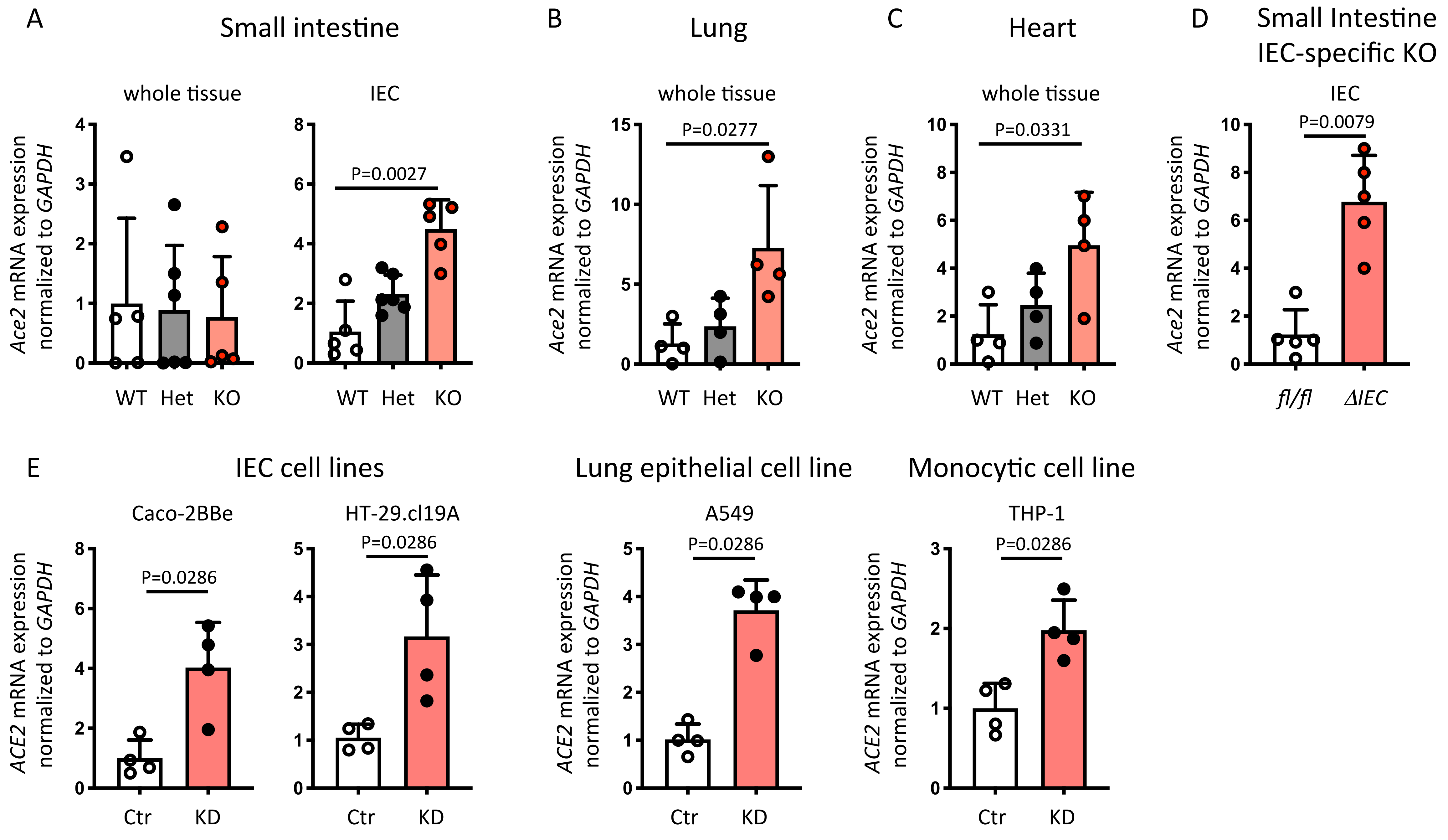


**Supplementary Figure 1. Loss of PTPN2 promotes ACE2 expression. A-D:** Intestinal epithelial cells (IEC) from the ileum, lung tissue, and heart tissue from 3-week-old wild-type (WT), *Ptpn2* heterozygous (Het) or *Ptpn2* knock-out (KO) mice (D), or 8-12 week old mice in which *Ptpn2* was specifically deleted in IECs (ΔIEC) or their control littermates (fl/fl) (E) was analyzed for *Ace2* mRNA expression*.* Data is normalized to *Gapdh* and the average of WT or fl/fl mice respectively. **E)** Caco-2BBe, HT-29.cl19A, A549, and THP-1 cells expressing non-targeting control (Ctr) or *PTPN2*-specific (KD) shRNA were analyzed for mRNA expression of *ACE2.* Data is normalized to *GAPDH* and the average of Ctr shRNA expressing cells. Statistical differences are indicated in the figures (One-way ANOVA (A-C) or Student's t-test (D+E); n = 4). Each dot represents a biological replicate.

SUPPLEMENTARY FIGURE 2

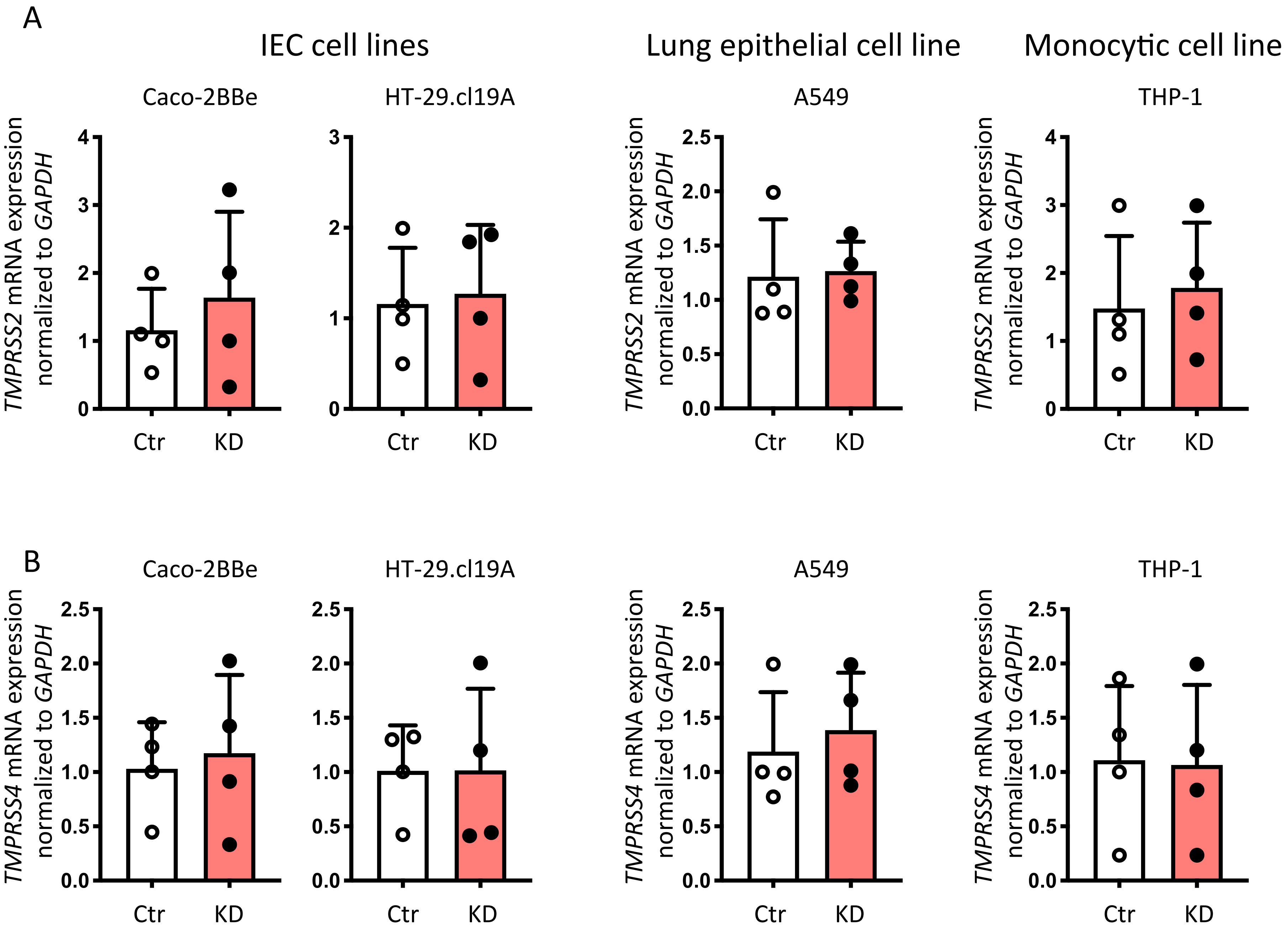


**Supplementary Figure 2. Loss of PTPN2 does not affect expression of TMPRSS2 and TMPRSS4.** Caco-2BBe, HT-29.cl19A, A549, and THP-1 cells expressing non-targeting control (Ctr) or *PTPN2*-specific (KD) shRNA were analyzed for mRNA expression of **A)** *TMPRSS2* and **B)** *TMPRSS4.* Data is normalized to *GAPDH* and the average of Ctr-shRNA-expressing cells. Statistical differences are indicated in the figures (Student's t-test; n = 4). Each dot represents the average of an independent experiment with 2-3 technical replicates.

SUPPLEMENTARY FIGURE 3

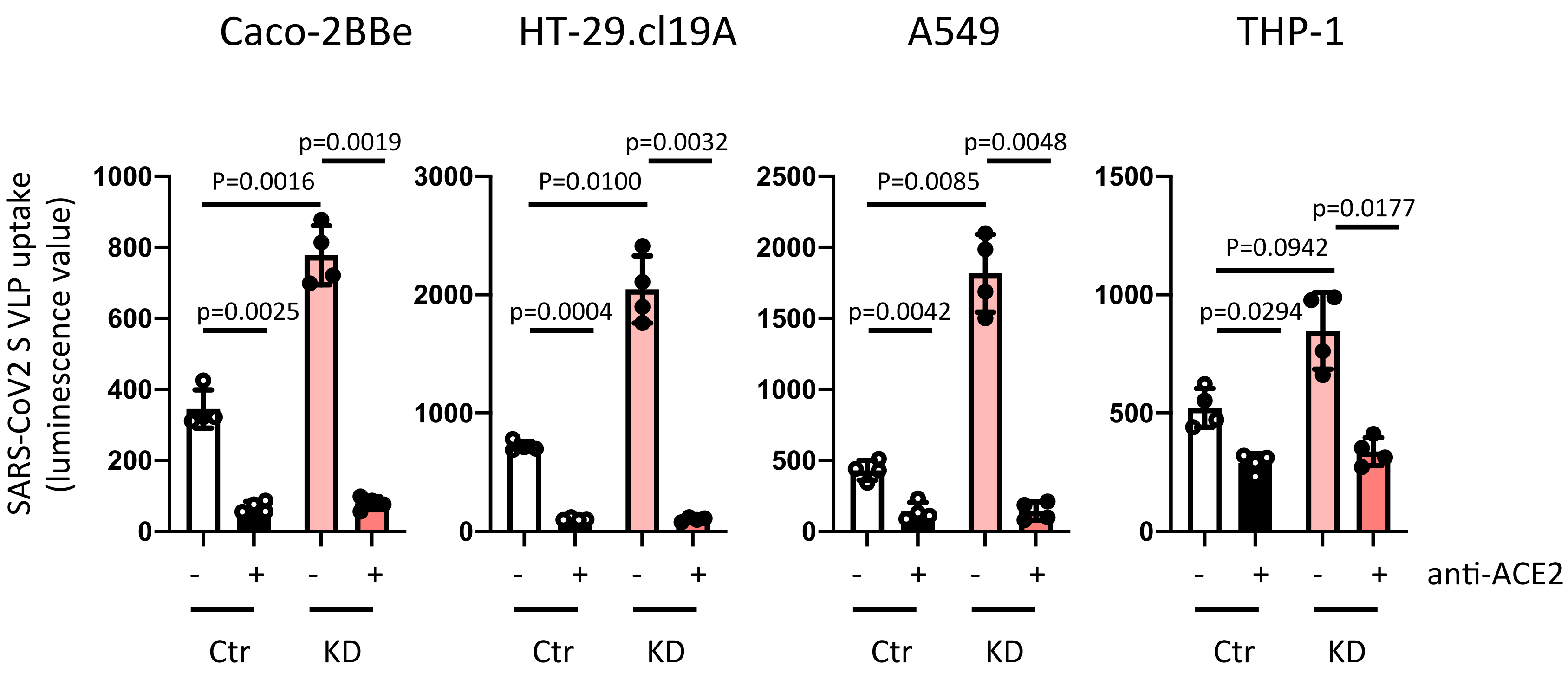


**Supplementary Figure 3. Increased uptake of SARS-CoV-2 spike expressing VLPs is dependent on ACE2.** Caco-2BBe, HT-29.cl19A, A549, and THP-1 cells expressing non-targeting control (Ctr) or *PTPN2*-specific (KD) shRNA were incubated with an inhibitory anti-ACE2 antibody prior to infection with VLPs expressing SARS-CoV-2 spike protein. 48 h after infection, luminescence was measured as an approximation of VLP uptake. Statistical differences are indicated in the figure (One-way ANOVA, n = 4). Each dot represents the average of an independent experiment with 2-3 technical replicates.

SUPPLEMENTARY FIGURE 4

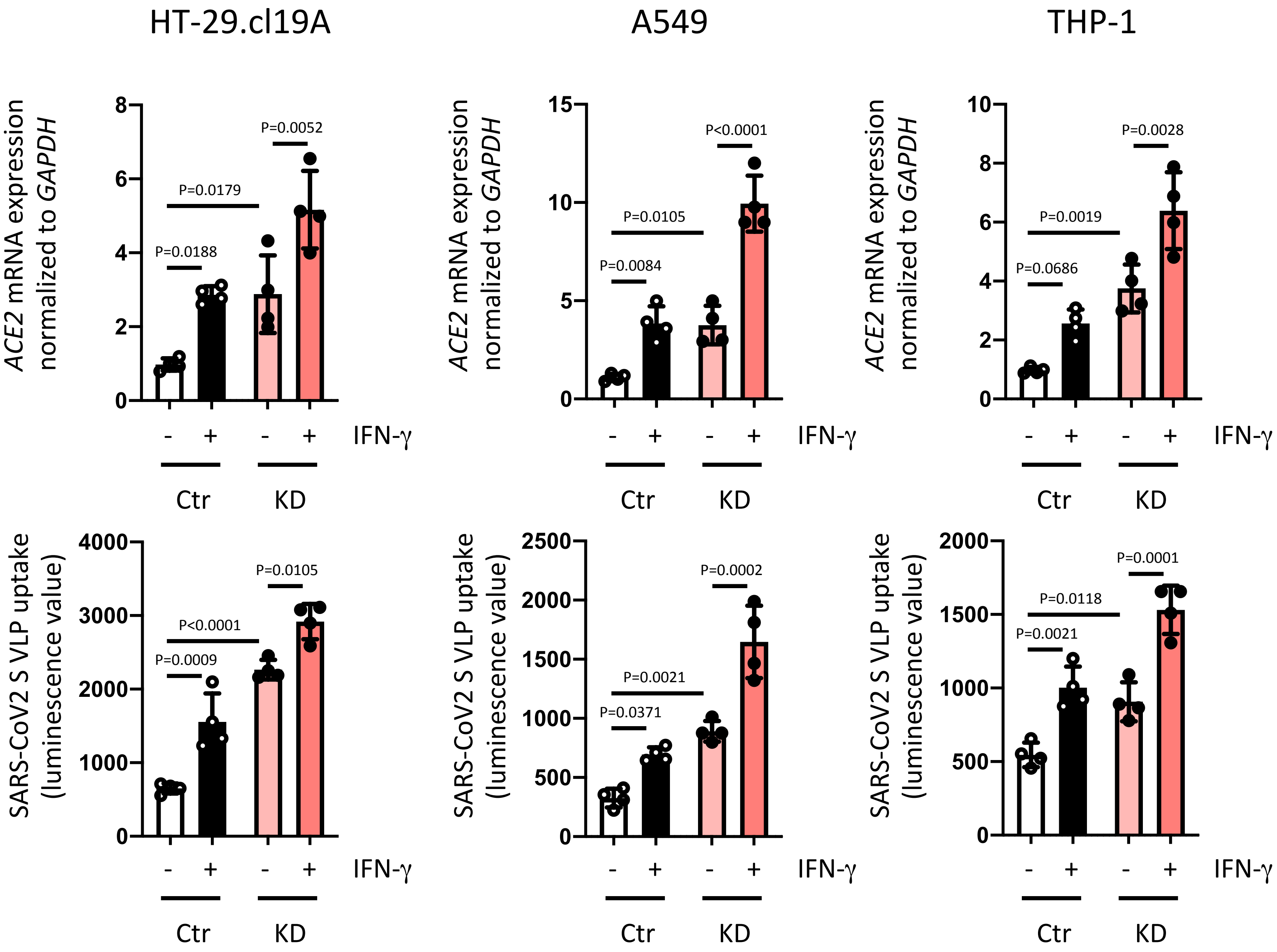


**Supplementary Figure 4. IFN-γ promotes uptake of SARS-CoV2 spike-expressing VLPs in several cell types.** HT-29.cl19A, A549, and THP-1 cells expressing non-targeting control (Ctr) or *PTPN2*-specific (KD) shRNA were infected with VLPs expressing SARS-CoV-2 spike protein in the presence or absence of IFN-γ. *ACE2* mRNA expression normalized to *ACTB* and untreated Ctr cells was measured after 24 h; Luminescence as an approximation of VLP uptake was measured after 48 h. Statistical differences are indicated in the figure (One-way ANOVA, n = 4). Each dot represents the average of an independent experiment with 2-3 technical replicates.

SUPPLEMENTARY FIGURE 5

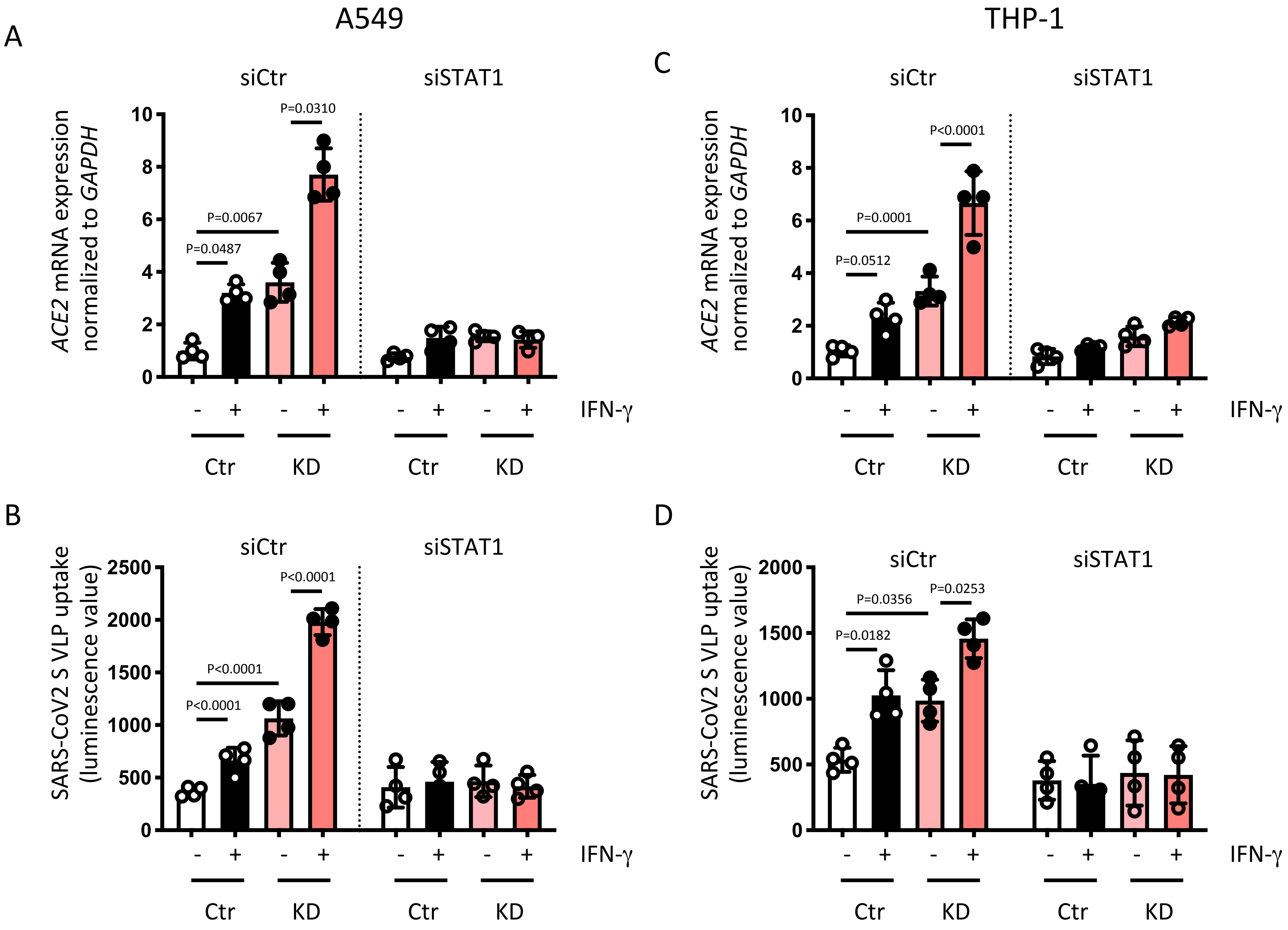


**Supplementary Figure 5. Silencing of STAT1 prevents IFN-γ and PTPN2-KD-mediated increase in ACE2 expression and uptake of SARS-CoV2 spike-expressing VLPs in several cell types.** HT-29.cl19A, A549, and THP-1 cells expressing non-targeting control (Ctr) or *PTPN2*-specific (KD) shRNA were treated with non-targeting control (siCtr) or STAT1-specific (siSTAT1) siRNA prior to infection with VLPs expressing SARS-CoV-2 spike S protein in the presence or absence of IFN-γ. **A+C)** for 24 h and analysis of *ACE2* mRNA expression and **B+D)** for 48 h and supernatant analyzed for luminescence as an approximation of VLP uptake. Statistical differences are indicated in the figure (One-way ANOVA, n = 4). Each dot represents the average of an independent experiment with 2-3 technical replicates.

SUPPLEMENTARY FIGURE 6

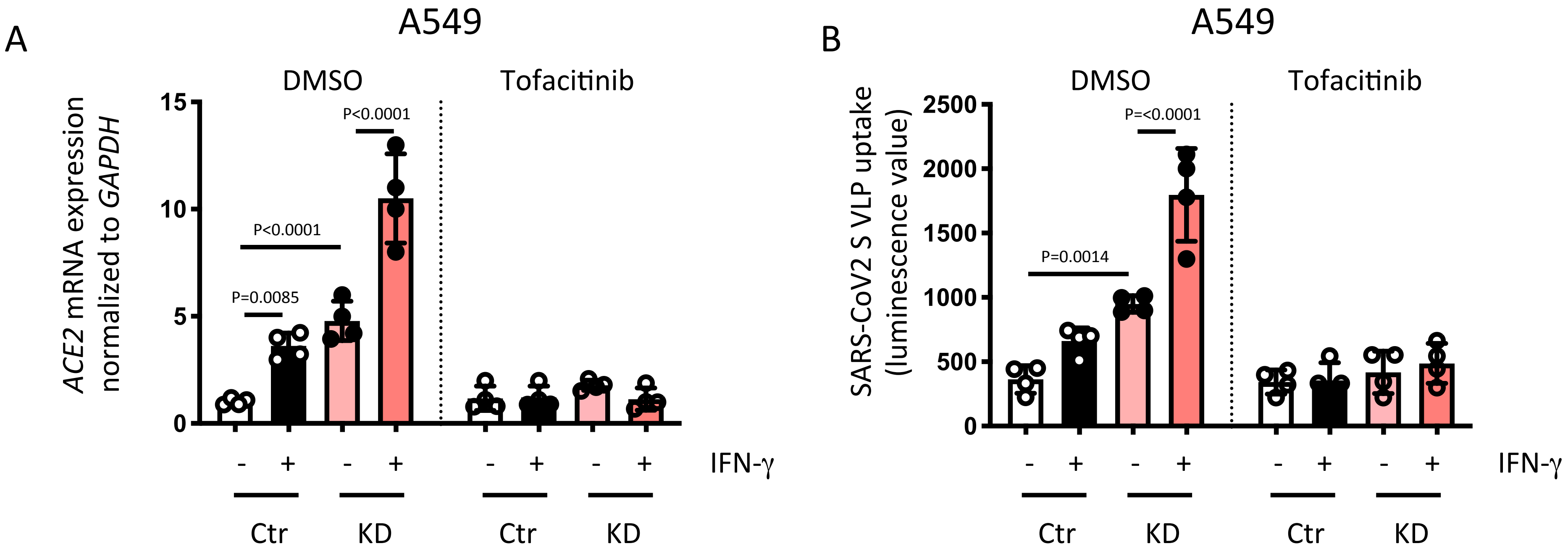


**Supplementary Figure 6. Tofacitinib treatment prevents elevated ACE2 expression and enhanced uptake of SARS-CoV2 spike S protein expressing VLPs in lung epithelial cells.** A549 cells expressing non-targeting control (Ctr) or PTPN2-specific (KD) shRNA were treated with vehicle (DMSO) or Tofacitinib for 1 h prior to infection with VLPs expressing SARS-CoV-2 spike protein in the presence or absence of IFN-γ. **A**) After 24 h ACE2 mRNA expression normalized to *GAPDH* and untreated control cells were measured. **B)** After 48 h the supernatant was analyzed for luminescence as an approximation of VLP uptake. Statistical differences are indicated in the figure (One-way ANOVA, n = 4). Each dot represents the average of an independent experiment with 2-3 technical replicates.
